# Disrupting dorsolateral prefrontal cortex by rTMS reduces the P300 based marker of deception

**DOI:** 10.1101/060723

**Authors:** Inga Karton, Anni-Bessie Kitt, Talis Bachmann

## Abstract

It is well known that electroencephalographic event related potential component P300 is sensitive to perception of critical items in a concealed information test. However, it is not known whether the relative level of expression of P300 as a neural marker of deception can be manipulated by means of non-invasive neuromodulation. Here, we show that while P300 exhibited systematic amplitude differences in response to the more as well as the less significant stimuli items encountered at the “crime scene” compared to neutral items, offline rTMS to dorsolateral prefrontal cortex attenuated P300 amplitude in response to the critical items. Yet, the individual subjects showed different sensitivity of the P300 as the marker of concealment. We conclude that rTMS can be used for subduing electrophysiological markers of deception, but this effect depends on whether the subject belongs to the group of CIT-sensitive individuals.

## 1. Introduction

The brain works differently when a person lies compared to how the brain works when this person tells the truth (Ganis et al., 2003; Langleben et al., 2005; Ganis and Keenan, 2009; Kozel et al., 2009; Jiang et al., 2013). This obvious fact makes it possible to develop objective methods of deception detection based on psychophysiology and brain imaging. In the various versions of the concealed information test (CIT) psychophysiological responses to critical (i.e., the so-called relevant probe, potentially incriminating) items are compared to the responses to neutral (i.e., the irrelevant, contextually not significant) items whilst subjects are trying to hide or deny that they have specific contextual knowledge of the critical items. If the critical items lead to enhanced responses compared to the responses to neutral items, possession of concealed information can be inferred.

Traditionally, CIT was used together with polygraph recordings, revealing enhanced respiratory and/or galvanic skin responses to critical items (e.g. Lykken, 1959, 1979; Ben-Shakhar and Elaad, 2003). However, in a more modern tradition the CIT is often combined with electroencephalography (EEG) to reveal deception-related event related potentials (ERPs). By now the relevance of the P300 component in connection with deception can be regarded as being well founded in research (Ambach et al., 2010; Rosenfeld and Labkovsky, 2010; Verschuere et al., 2011). If a deception-related critical (probe) stimulus is presented, the P300 in response to this stimulus is enhanced compared to irrelevant stimuli (Ambach et al., 2010; Rosenfeld et al., 2013).

However, the above mentioned electrophysiological measures of deception are correlational if viewed from the methodological point of view – brain-imaging markers correlate with certain behavioral processes, but causal effects cannot be definitely revealed. A somewhat different tradition of neurobiological research on deception combines brain imaging with non-invasive brain stimulation (for reviews see Rogasch and Fitzgerald, 2013 or Shafi et al., 2012, for example). This approach is capable of examining causal effects and therefore increase methodological rigor of the studies of brain mechanisms of deception (Gamer et al., 2007; Priori et al., 2008; Karton and Bachmann, 2011; Karton et al., 2014a, 2014b; Luber et al., 2009; Mameli et al., 2010). Despite this potential, there have been no studies examining the effects of brain stimulation on deception-related P300 ERP responses so far.

Several previous studies have reported that the dorsolateral prefrontal cortex (DLPFC) is involved in deceptive behavior (Priori et al., 2008; Christ et al., 2009; Mameli et al., 2010; Ito et al., 2012). In one of our earlier studies (Karton and Bachmann, 2011) we explored the causal effect of manipulation of DLPFC on deception related behavior and found that repetitive offline transcranial magnetic stimulation (rTMS) of the right-hemisphere DLPFC relatively decreased untruthful responses, whereas left-hemisphere DLPFC stimulation relatively increased lying. Because a change in the amplitude of ERP/P300 is the best known brain-potential signature of deception in the concealed information detection test and because TMS has been shown to affect P300 (Hansenne et al., 2004; Torii et al., 2012), it would be important to know whether rTMS targeted at DLPFC has any effect on the extent of expression of P300 in the context of ERP based CIT. If this kind of effect will be found we might also ask are there any hemispheric differences analogous to what was found in the behavioral spontaneous lying study. This is important both for theoretical analysis of the brain mechanisms involved in the subjects’ behavior in the CIT-like tasks and for practical purposes when manipulation with subjects’ sensitivity to critical stimuli operationalized by deception-related ERPs might be desirable. If rTMS can lead to higher sensitivity of ERPs to deception, the ERP based deception detection methods can be improved.

Many of the above mentioned psychophysiological measures are able to reflect the deception related significance of an item quite robustly when combined with CIT. It is worth keeping in mind, however, that the CIT was first envisioned as a test for recognition not deception (Rosenfeld, 2011). Therefore, it is not surprising that sensitivity of deception detection tests varies depending on the exact specifics of the experimental paradigm (Winograd and Rosenfeld, 2014). For example, results depend on the time delay between the crime episode and the administration of a CIT (Carmel et al., 2003), or on the types of questions that the subjects are confronted with (Carmel et al., 2003; Jokinen et al., 2006; Ambach et al., 2011). Characteristics of the concealed information/knowledge are another likely source of variability. The significance of crime related knowledge is one such source. Ambach et al. (2011) compared two different questioning formats (“did you see?” vs “did you steal?”) with two different encodings of items of interest (only seen vs seen and stolen items). Thus, the level of significance was different for these two types of items and depending on the item type only one question required deception. They found that both types, irrespective of the question, elicited different physiological responses (skin conductance, electrocardiogram, respiration, and finger pulse) compared to completely new items. Importantly, responses to the actually stolen objects were also different from responses to the merely seen objects. This allows distinction between markers of “guilty knowledge” and “not guilty knowledge”.

By gaining support from the work of Ambach et al. (2011) it is important to further understand how levels of information significance influence the sensitivity of deception detection with a CIT and to examine whether the level of expression of the corresponding ERP-markers of deception can be manipulated by brain stimulation. The combination of CIT with EEG, supplemented by rTMS should be particularly well suited for pursuing this task. This combination allows a superior temporal resolution of the evoked neural processes in response to a crime related item (and its significance), precise targeting of the brain areas likely involved in deception and studying causal effects in addition to the purely correlational brain-imaging data. It is important to test how rTMS targeted at DLPFC affects the P300 response as a function of stimulus significance. After all, it may be possible that the potential effects are mediated through recognition processes and not through guilty knowledge *per se*.

In the current study we used a variety of a CIT protocol with the mock crime scenario as follows. We distinguish between three types of stimuli: the critical stimulus (stolen by the subject), familiar stimuli (present during the enactment of the crime) and neutral stimuli (no prior exposure). The goal of the experiment was to investigate how rTMS targeted at DLPFC affects P300 as a function of stimulus significance in the CIT context. As previous research has shown that rTMS to DLPFC affects truthfulness of behavioral responses (Karton and Bachmann, 2011; Karton et al., 2014a; Karton et al., 2014b) and as P300 has been found to be susceptible to rTMS effects in the context of cognitive control (Hansenne et al. 2004; Torii et al., 2012), it is advisable to test whether P300 based markers of deception are susceptible to rTMS effects in the CIT context. This constitutes the main aim of our study.

To state the working hypotheses we need also to consider specific information related to DLPFC-targeted rTMS effects on deceptive behavior. On the one hand it appears that especially right-hemisphere rTMS targeted at DLPFC influences deceptive behavior (Karton and Bachmann, 2011; Karton et al., 2014a) and thus it is expected that rTMS has a significant effect on P300 based markers of deception. On the other hand, clear disruptive rTMS effects on P300 have been found specifically with left-hemisphere rTMS of DLPFC (Torii et al., 2012). Thus, in order to have a clearer picture of the putative rTMS effects on P300 based deception markers in the CIT context, DLPFC of both hemispheres will be stimulated. We hypothesize that right but not left DLPFC rTMS will have an effect on the P300 difference between the conditions of neutral and critical stimulus presentation whereas left DLPFC rTMS will change P300 parameters uniformly regardless of the stimulus type. For the behavioral task we employed sheets of paper with words indicating goods to steal. The subjects were instructed to imagine stealing one of these items from a store when they choose the label for that particular item. Words were used as stimuli during the CIT as well.

## 2. Method

### 2.1 Participants

All subjects who participated in our experiment were healthy and had normal or corrected to normal vision. They gave written informed consent before participation. The experiments were approved by the Research Ethics Committee of the University of Tartu and were conducted according to the principles set in the Declaration of Helsinki. Some of the subjects received monetary compensation for participation, others were awarded partial course credits.

25 subjects (5 male, 20 female) participated in the experiment. The data of two male and five female subjects were excluded due to noisy EEG recordings and extensive blink artifacts. The age of the remaining 18 subjects ranged from 20 to 40 years (mean age 25.18 years, standard deviation (SD) 5.63 years). Subjects were randomly divided into two stimulation groups: 9 subjects (1 male and 8 female) received rTMS (repetitive transcranial magnetic stimulation) and sham stimulation targeted to left DLPFC (dorsolateral prefrontal cortex) and 9 subjects (2 male and 7 female) received rTMS and sham stimulation targeted to right DLPFC. (In the sham condition, the coil was pressed perpendicularly to the subject’s head and no real magnetic pulses were generated.)

### 2.2 Experimental procedures

Our experimental task was analogous to other variations of the Guilty Knowledge Test (GKT) and the Concealed Information Test (CIT). The experiment started with the mock crime scenario (a “shoplifting” episode). We used cards with words referring to five kinds of items ‘easy’ to steal (e.g., chewing gum, candy, fruit, etc.). In each session, three words from five possible word alternatives written on cards were selected at random and put face up on a table next door. The subjects were instructed to enter that room and imagine that they are in the supermarket about to steal something. For this, one of the three cards had to be taken. Subjects had to write the name of the item on the opposite side of the card and specify it more precisely, e.g. by naming some favorite brand. Then the card had to be placed into a folder as a “shopping bag” whereupon subjects returned to the room of the experiment, bringing the folder with them. Subjects were told that the purpose of the experiment is to discover “stealing” using EEG recordings and they should hide their “crime” related knowledge.

The five possible word alternatives belonged to three categories of stimuli for our experimental task. One was the item actually stolen by the subject (critical stimulus category). Two additional items that were also present during the shoplifting episode, but not stolen belonged to the category of familiar stimuli. Two word alternatives were completely new items to be shown only later at the CIT task stage; these words were previously unseen by the subject (neutral stimuli category). After the shoplifting episode (experiment step one), the EEG cap was fitted to the subject’s head (experiment step two), followed by blocks of sham/rTMS (experiment step three) and the CIT stimuli presentation with concurrent EEG recordings (experiment step four).

Subjects were seated at a distance of 70 cm from the computer monitor (Eizo FlexScan T550, 1024 × 768 pixels, 85 Hz refresh rate). During the experimental step four word stimuli were presented foveally on a computer screen in random order. Each stimulus was presented 18 times per block. Thus, there were 90 stimuli in each block. The words were printed with high-contrast dark letters on a light background. The luminance of the background was 80 cd/m^2^. Each trial begun with a fixation cross in the middle of the screen. Participants were instructed to fixate the cross and refrain from any kind of movement (e.g. blinking, turning their head etc.) until the onset of the response screen. After 1176 ms the fixation cross was replaced by the word stimulus for 400 ms, followed again by the fixation cross for 1094 ms. Then, a question appeared for 1000 ms, followed by a yes/no response screen. One of the two possible questions appeared at random: “Was this word written on one of the cards?” or “Is the card with this word held by you?”. In the first case subjects should answer honestly, in the second case they should deny having the card. (Throughout the experiment, subjects generally adhered to the instructions and made few mistakes of response – see part 3.1 further on.) Note, however, that in the present study we restrict our analyses to ERPs in response to the different types of stimuli and not to the two questions. Responses were given on a standard computer keyboard. After responding, subjects initiated the next trial by pressing the space bar.

Each block of the experimental task was associated with “off-line” rTMS in order to capitalize on earlier research showing suitability of this format in order to have an inhibitory effect on the cortical areas involved in deception (Hallett, 2007; Luber et al., 2009; Karton and Bachmann, 2011). “Off-line” protocol means that rTMS was not applied during the block of trials with stimuli presentation, but only before this block begun. One group of subjects received rTMS and sham stimulation targeted to left DLPFC. Another group of subjects received rTMS and sham stimulation targeted to right DLPFC. Each subject participated in two experimental sessions, carried out on different days. Each session comprised a different sequence of four experimental blocks: AA/BB or BB/AA, where A is sham stimulation and B is real rTMS. The order of the two sequences was counterbalanced between subjects within both groups of subjects.

Prior to each experimental block a train of 1-Hz rTMS (360 pulses over the course of 6 min) or equivalent sham stimulation was delivered either to the left or to the right DLPFC. At this point it is necessary to note that little is known about the duration of “off-line” stimulation effects in PFC. Hansenne et al. (2004) maintained that 1-Hz rTMS produces inhibitory effect only when the duration of the stimulation is about 15 min and according to Robertson and colleagues (Robertson et al., 2003) and Thut and Pascual-Leone (2010) the effect of stimulation in DLFPC is diminishing after 5-10 (15) minutes from the end of stimulation. According to Eisenegger and colleagues (Eisenegger et al., 2008) the prefrontal rTMS causes increase in rCBF under the stimulation site which lasts about 9 minutes. In order to collect enough data for EEG analysis so that there is a sufficient number of trials corresponding to the still present rTMS effect the need to split the experiment between two days for each participant was therefore acknowledged.

MRI-assisted NBS (Navigated Brain Stimulation; Nexstim Ltd., Helsinki, Finland) with a figure-of-eight coil was used for stimulation. To mask the coil-generated clicks and to reduce differences between real rTMS and sham stimulation, music was played over earphones in both cases. For sham stimulation, however, the music was mixed with an audio recording of the TMS clicks while the coil was pressed perpendicularly to the subject’s head and no real magnetic pulses were generated. Stimulation (either sham or real) was immediately followed by the experimental task.

Intensities close to the motor threshold (MT) are typically used as a guide for the stimulus intensity needed for prefrontally applied TMS (Kähkönen et al., 2004). In our experiment, stimulation intensity was set at 80% of the individual MT (measured as a barely noticeable twitch of the thumb). The stimulation intensity used for different subjects according to the above mentioned 80% MT criterion ranged between 29% and 42% of maximal stimulator output.

### 2.3 EEG and data analysis

We used the Nexstim eXimia EEG-system with 60 carbon electrodes cap (Nexstim Ltd., Helsinki, Finland) for EEG recordings. The impedance at all electrodes was kept below 10 kΩ. The EEG signals were referenced to a reference electrode placed on the forehead and sampled at 1450 Hz. All signals were amplified with a gain of 2000 and filtered with a hardware based bandpass filter of 0.1–350 Hz. The vertical electro-oculogram (VEOG) was recorded via two additional electrodes placed above and below the participants’ left eye. All recorded EEG data were analyzed with Fieldtrip (http://fieldtrip.fcdonders.nl; version 14-12-2013), an open-source MATLAB toolbox.

Bioelectrical activity was recorded from 15 electrodes: frontal (electrodes AF1, F1, F5, AF2, F2, F6), parietal (electrodes P3, PO3, P4, PO4), temporal (electrodes TP7, TP8), and central (electrodes C3, C4, CZ). After the initial recording, data were low-pass filtered (zero phase shift Butterworth filter 30 Hz) and segmented into trials from -200 ms to 1000 ms relative to stimulus onset. The data were manually checked for any artifacts, including eye movements and blinks. All trials contaminated by artifacts were discarded from further analysis. Data were baseline-corrected with a 100 ms window prior to the stimulus onset. For cleaner ERP traces in figures, data was filtered with a 10-Hz low-pass filter instead of the 30 Hz low-pass filter used for data analysis.

Average ERP potentials were computed for each subject in each condition and for each electrode. The activity from single electrodes was pooled into two regions of interest (ROIs). The ROIs were frontal (electrodes AF1, F1, F5, AF2, F2, F6), and parietal (electrodes P3, PO3, P4, PO4). We ended up with these two ROIs because of the TMS related constraints and due to the actual sensitivities of the channels in each experiment.

The experimental conditions specified according to stimulus category types were neutral (stimuli that were not encountered previously during the mock crime), familiar (stimuli that were encountered during the mock crime, but not stolen) and critical (the stolen stimulus). On average, the following number of trials were available: for the neutral condition: mean = 111.3, SD = 22.9; for the familiar condition: mean = 110.9, SD = 23.3; for the critical condition: mean = 56.0, SD = 12.1. The number of available trials was very similar for TMS and SHAM conditions.

Peak-to-peak amplitude was used for the analysis of P300 (as recommended by Soskins et al., 2001). The algorithm first identified the 100 ms long segment between 300 and 800 ms after stimulus onset which had the highest positive average amplitude. P300 latency is defined as the midpoint of this segment. Next, the algorithm identified the 100 ms long segment between P300 latency and 1000 ms after stimulus onset, which had the highest negative average amplitude. P300 peak-to-peak amplitude is defined as the difference between the highest positive and the highest negative average amplitude. Note that for this procedure data was first averaged over the electrodes within each electrode group. Note also, that the algorithm was applied on each individual ERP.

### 2.4 Statistical analysis

Statistical analyses were performed with R (version 3.0.3), a freely available and powerful statistical programming language. Repeated-measures analysis of variance (ANOVA) was used to assess the effects of our experimental conditions. If the sphericity assumption was violated according to Mauchly’s test for sphericity, p-values were corrected with the Greenhouse-Geisser method. Only the corrected p-values are reported. As recommended by Bakeman (2005), generalized eta-squared is used to report effect sizes of our ANOVA results. Planned comparisons and post-hoc contrasts were carried out via dependent samples t-test. Post-hoc contrasts were corrected with the Holm– Bonferroni method. Unless indicated otherwise, only the corrected p-values are reported. Cohen’s d is reported as an estimate of effect size for the dependent samples t-tests.

To anticipate our results, we replicated previous findings by observing a significantly higher mean P300 amplitude for the critical compared to the neutral condition. To assess the reliability of this experimental effect within each participant, we performed a bootstrapping test as recommended by Rosenfeld, Biroschak, and Furedy (2006). For each subject, 100 iterations of the standard ERP extraction procedure were performed. On each iteration, 25 trials were randomly chosen from all available trials in the critical and in the neutral condition. These trials were averaged according to their condition and the peak-to-peak P300 amplitude was calculated using the above described criteria. Finally, the difference in P300 amplitude for the critical and the neutral condition on a given iteration was computed. Thus, after a hundred iterations we obtain for each subject a percentage of iterations where the P300 amplitude of the critical condition is higher than the P300 amplitude of the neutral condition. Or put differently, we obtain a percentage of iterations where the previously found experimental effect is evident for a given subject.

## 3. Results

### 3.1 Behavioral results

Remember that on each trial participants responded to one of two possible questions. The first question was “Was this word written on one of the cards?“. Participants were instructed to answer truthfully when this question appeared. The overall error rate is quite low for this question, albeit with a few outlier participants. On average, participants denied having seen the critical item on 3.6% of the trials (median = 1.4%, SD = 7.7%, range = 0 – 33.3 %). Participants denied having seen the familiar items on 4.8% of the trials, on average (median = 2.1%, SD = 8.3%, range = 0 – 34.5 %) and reported having seen the neutral items on 8.5% of the trials, on average (median = 1%, SD = 23%, range = 0 – 94.8 %).

The second question was “Is the card with this word held by you?” and participants had to deny possession of any cards. The overall error rate is quite low for the second question as well, but there was one outlier participant. On average, participants admitted to possessing the critical item on 1.1% of the trials (median = 0%, SD = 3.5%, range = 0 – 14.8 %). Participants reported the possession of familiar items on 1.1% of the trials, on average (median = 0%, SD = 3.6%, range = 0 – 15.6 %) and the possession of neutral items on 0.2% of the trials, on average (median = 0%, SD = 0.4%, range = 0 – 1.4 %).

Considering the high variability in error rates it is likely that a few subjects understood the task incorrectly and/or made deliberate errors, thus sometimes giving incorrect answers. However, as we are interested in the EEG responses to the stimuli and not the questions and because stimuli always preceded questions by a considerable time interval we decided not to exclude these participants from the analyses.

### 3.2 EEG-TMS results

First, a four-way repeated-measures ANOVA with the factors electrode group (frontal and parietal), stimulus type (critical, familiar, neutral) and stimulation type (TMS and SHAM) as within-subject factors and stimulation side (left and right) as a between-subjects factor was performed to assess differences in P300 amplitude. The ERP waveforms per condition can be seen in figure 1. The main effect of electrode group was significant (F(1,16) = 58, p = 1.0e-06; ηG^2^ = 0.45). This was due to the much lower peak-to-peak amplitude of frontal electrodes (m = 4.5, SD = 1.9) compared to parietal electrodes (m = 10.1, SD = 3.8) (see table 1). The main effects for stimulation condition and stimulation side were not significant (F(1,16) = 1.1, p = 0.32; ηG^2^ = 0.002 and F(1,16) < 1.0, respectively). There was a near significant trend for the main effect of stimulus type (F(2,32) = 3.0, p = 0.07; ηG^2^ = 0.009), but there was a significant interaction between stimulus type and stimulation type (F(2,32) = 3.6, p = 0.04; ηG^2^ = 0.007). All other interactions were not significant (all F’s < 1.9, all p’s > 0.19, all ηG^2^ < 0.004).

**Figure 1.**
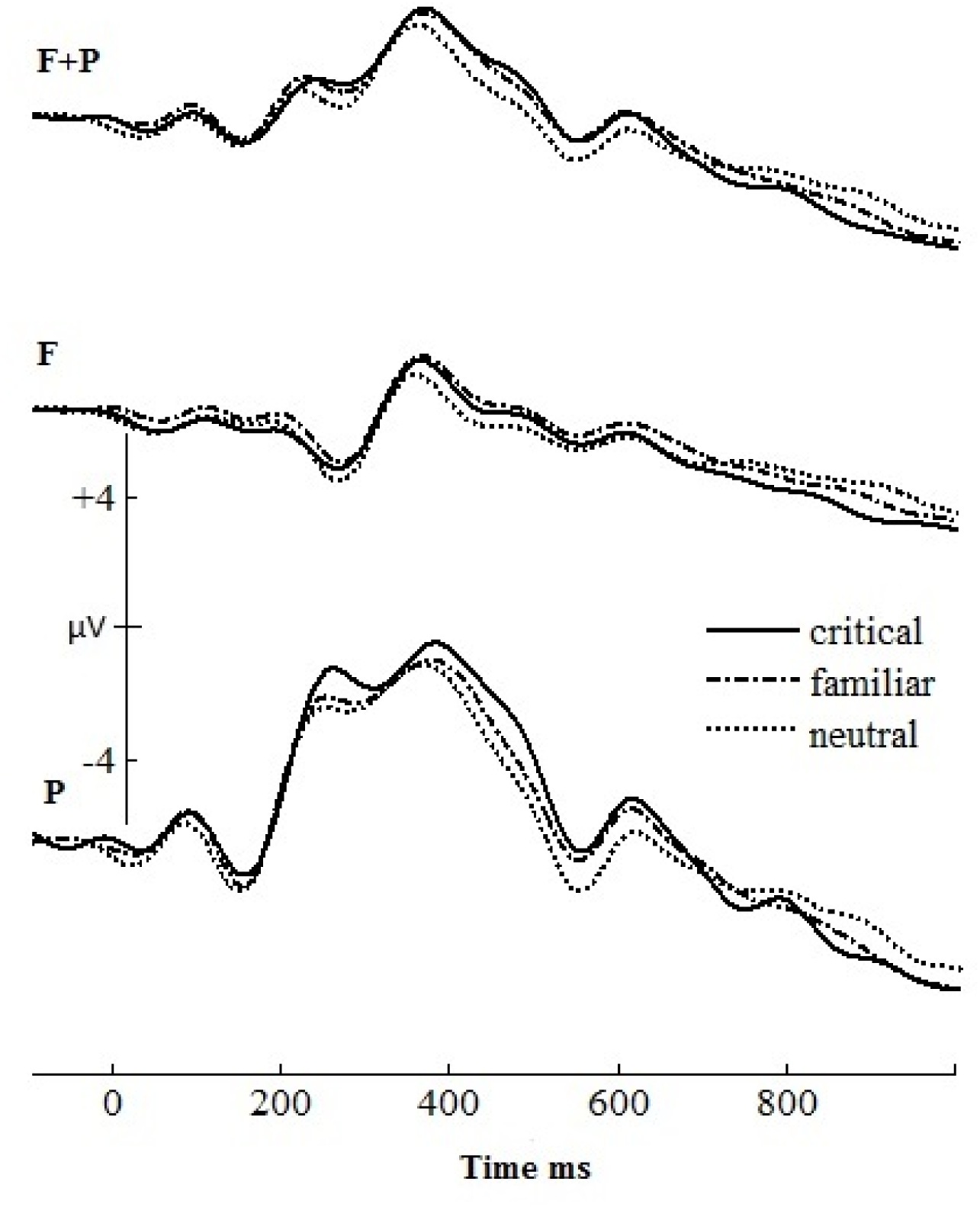
ERPs of the critical, familiar and neutral conditions after SHAM stimulation recorded from frontal (F) and parietal electrodes (P). To observe the absolute values of the voltage scale for each ERP respective baselines should be rescaled to voltage zero.

**Table 1.** Paired t-test means and standard deviations, averaged over the P300 at frontal and parietal electrodes. Notes: N = 18; ^a^SHAM Neutral < SHAM Critical, p < 0.001; TMS Critical < SHAM Critical, p < 0.05.

| Condition | Critical/probe |  | Familiar |  | Neutral/Irrelevant |  |
| --- | --- | --- | --- | --- | --- | --- |
|  | mean | SD | mean | SD | mean | SD |
| SHAM | 8,14<sup>a,b</sup> | 2,46 | 7,451 | 2,76 | 6,77<sup>a</sup> | 2,87 |
| TMS | 7,22<sup>b</sup> | 3,14 | 7,24 | 2,94 | 7,13 | 2,59 |

Planned comparisons were carried out to investigate which conditions differed significantly from each other in terms of the P300 amplitude they elicited. Because no significant interactions between electrode groups, stimulus type and/or stimulation side were found paired t-tests are carried out on the average P300 amplitudes over both electrode groups and both stimulation sides. First, we investigated which stimulus types elicit differing P300 amplitudes in response to TMS versus SHAM stimulation. There was a significant difference between the conditions of critical stimuli after TMS and after SHAM stimulation (t(17) = -2.3, p = 0.04, d = 0.53). The differences in P300 between the conditions of familiar stimuli after TMS vs after SHAM (t(17) = -0.5, p = 0.63, d = 0.12) and the conditions of neutral stimuli after TMS vs after SHAM (t(17) = 1.2, p = 0.25, d = 0.28) were not significant.

Second, we investigated if P300 recorded in response to stimulus types are different within stimulation conditions. Whereas P300 in the conditions of the three stimulus types did not differ from each other in the TMS condition (critical vs neutral: t(17) = 0.2, p = 0.84, d = 0.05; critical vs familiar: t(17) = -0.06, p = 0.96, d = 0.01; familiar vs neutral: t(17) = 0.31, p = 0.76, d = 0.07), in the SHAM condition this difference was significant when P300 was compared between the conditions of critical stimuli and neutral stimuli (t(17) = 4.6, p = 0.0002, d = 1.1). The comparisons between the conditions of critical and familiar (t(17) = 1.8, p = 0.09, d = 0.42) and familiar and neutral stimuli (t(17) = 1.7, p = 0.11, d = 0.4) were not significant in the SHAM condition. Figure 2 and table 1 show the P300 amplitudes and significant differences between our experimental conditions.

**Figure 2.**
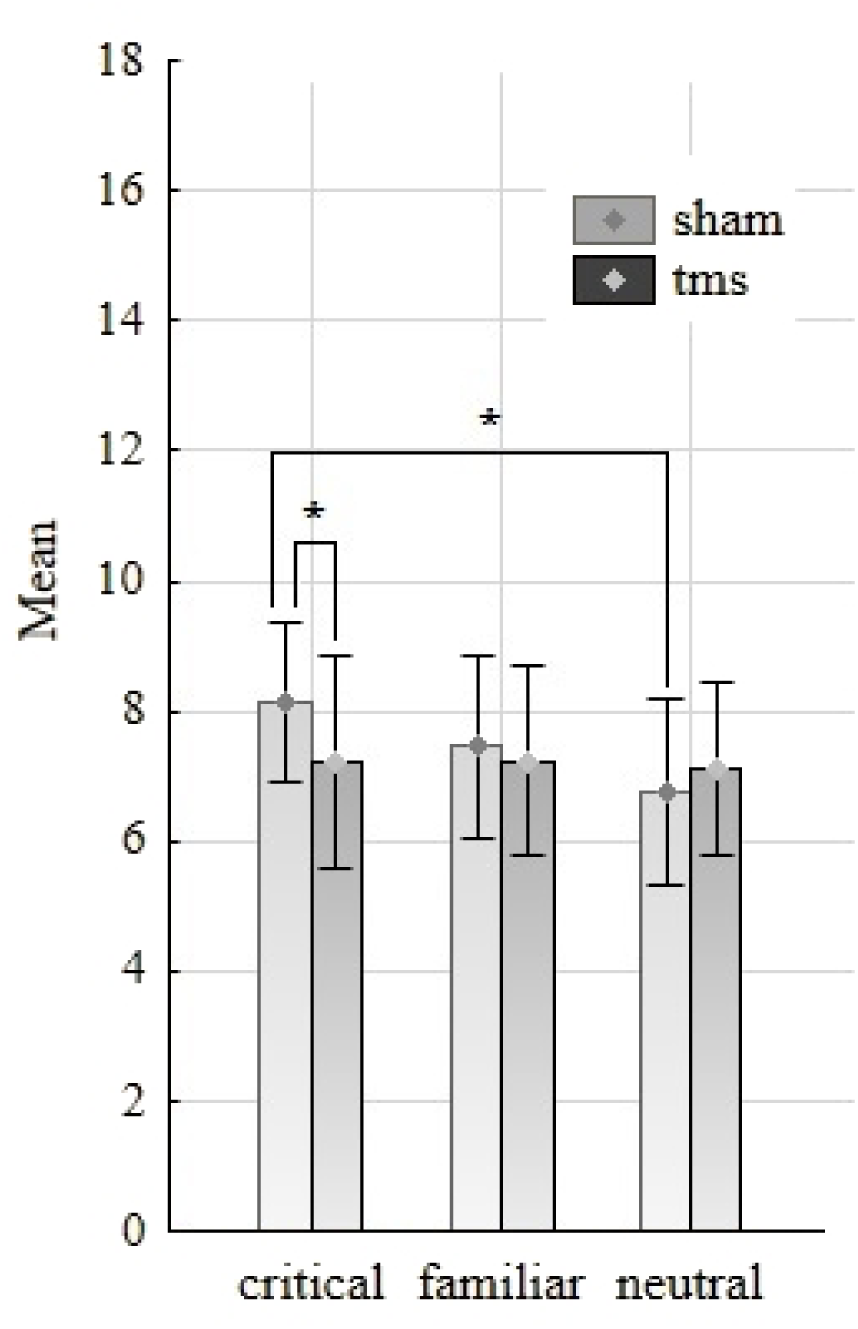
Mean peak-to-peak amplitudes (μV) and standard errors for the critical, familiar and neutral stimulus conditions per stimulation condition (SHAM/TMS). Amplitudes are averaged over frontal and parietal electrode groups (paired t-test differences, *p<0.05).

Thus, the results show that the P300 response to critical stimuli has higher amplitude if compared with the P300 response to neutral stimuli. This effect was abolished, however, if DLPFC was inhibited with rTMS prior to stimulus presentation. Interestingly, the P300 response to critical items exhibits decreased amplitude after rTMS to both left- and right-hemisphere DLPFC.

### 3.3 Intraindividual reliability

Let us now turn to the question about how reliable the differences for P300 amplitude in response to critical and neutral stimuli are within single subjects. After all, if P300 amplitude really indexes deceptive behavior universally, it should be evident and quantitatively confirmable within each participant. To investigate this matter the bootstrap method (Rosenfeld et al., 2006; see Methods for more details) was used to determine whether the critical-neutral differences were consistent enough within single subjects to detect deception.

First, the bootstrap test was conducted in the SHAM condition, independent of stimulation side. Table 2 shows the results of all the 18 subjects. For each subject the percentage of iterations where the P300 amplitude of the critical condition exceeded the P300 amplitude of the neutral condition is shown. We will concentrate on the results from those bootstrap tests which were carried out over the average of frontal and parietal electrodes. First, these tests are in accordance with the results from the above described group analyses, where no interactions between electrode groups and other experimental factors were found. Second, these tests best discriminate between the critical and the neutral conditions. However, bootstrap tests over the parietal electrode group only are also reported for comparison. Evidently, only four participants (22%) reached the necessary 90% criterion for highly reliable deception detection. Figure 3 gives some examples of the ERPs of subjects with the highest and with the lowest hit rates on the bootstrap test.

**Figure 3.**
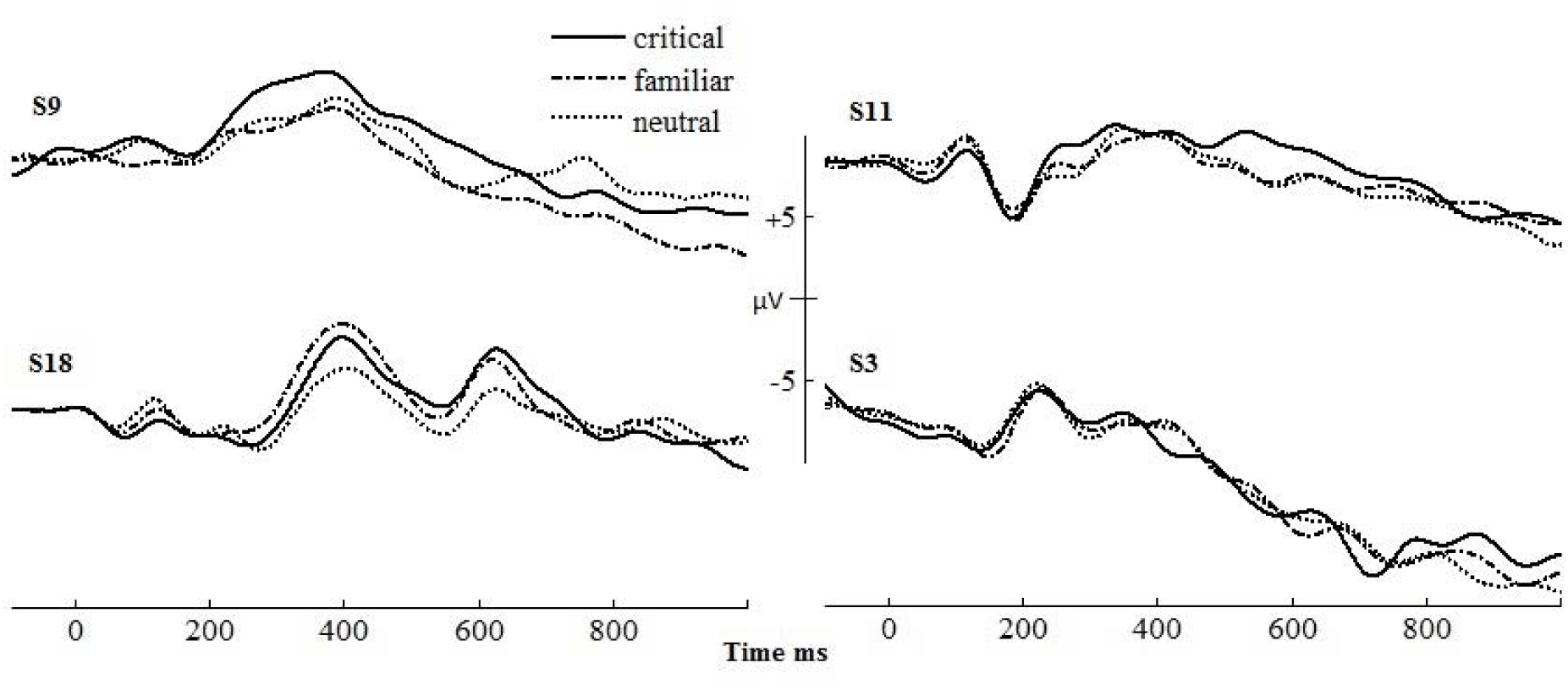
ERPs of individual subjects with high critical > neutral index (S9, S18) and low critical > neutral index (S11, S3). To observe the absolute values of the voltage scale for each ERP respective baselines should be rescaled to voltage zero.

**Table 2.** Bootstrap results. Note: Percent of iterations for which the critical condition was higher than the neutral condition in terms of P300 amplitude. In the first case only P300 amplitude of the parietal electrode group was used for the bootstrap tests. In the second case the mean over both electrode groups was used.

| Subject | Parietal | F+ P |
| --- | --- | --- |
| 1 | 72 | 80 |
| 2 | 76 | 74 |
| 3 | 67 | 74 |
| 4 | 84 | 85 |
| 5 | 98 | 96 |
| 6 | 29 | 59 |
| 7 | 79 | 95 |
| 8 | 58 | 55 |
| 9 | 86 | 96 |
| 10 | 81 | 74 |
| 11 | 29 | 36 |
| 12 | 43 | 52 |
| 13 | 22 | 24 |
| 14 | 83 | 80 |
| 15 | 63 | 89 |
| 16 | 34 | 68 |
| 17 | 54 | 55 |
| 18 | 93 | 91 |
| Mean % | 64 | 71 |
| Corr class 90% | 2 (11%) | 4 (22%) |
| Corr class 80% | 6 (33%) | 8 (44%) |

## 4. Discussion

The present study aimed at investigating whether non-invasive brain stimulation by offline rTMS has an impact on the relative expression of the best known ERP marker of deception, the augmented P300 to critical stimuli. We carried out this study in the context of a variant of a CIT protocol with significance of mock crime related items varied and also observed the effects at the individual subjects level. We distinguished between three categories of stimuli (critical, familiar and neutral) and expected to find differences between the critical and the neutral stimuli. By using rTMS, functionality of the brain area DLPFC (and possibly sites associated with it) known to be involved in deception was disrupted. This intervention was motivated by the results of our earlier studies (Karton and Bachmann, 2011; Karton et al., 2014a) showing that direct or indirect stimulation especially of the right-hemisphere DLPFC has an effect on deceptive responses. The results showed that (i) amplitude of P300 to critical items as compared to familiar and neutral items, was strongly reduced (effectively eliminated) and (ii) expression of the P300 effects varies considerably between individual subjects. P300 as a neural marker of deception was reliable in part of the subjects and unreliable with other subjects in the present study. Consequently, rTMS as a means to reduce sensitivity of ERP markers to deception in the CIT paradigm seems to be applicable only for a part of population. As the number of critical trials was relatively small in our experiment and as we used only one specific variety of the CIT paradigm it is possible that the proportion of population susceptible for the P300 based valid and sensitive lie-detection test may be larger (e.g., Rosenfeld et al., 2013) than was the case in this investigation.

It is known that right DLPFC is involved in cognitive control, avoidance and behavioral inhibition (Cho et al., 2012; Knoch and Fehr, 2007; Ott et al., 2011; Shackman et al., 2009; Tassy et al., 2012) while left DLPFC participates in reality monitoring, approach motivation / aggression, strategic behavior, naming and execution (Berkman and Lieberman, 2010; Fertonani et al., 2010; Hortensius et al., 2012; Huffmeijer et al., 2012; Ito et al., 2012; Ott et al., 2011; Steinbeis et al., 2012). Deception requires considerable cognitive control to inhibit habitual and reality-corresponding behavior, thus we expected that right DLPFC stimulation by 1-Hz rTMS will decrease P300 in response to critical stimuli relative to neutral stimuli compared to the control condition when sham stimulation is used.

We found that in the sham condition the P300 component exhibited systematic amplitude differences in response to critical items compared to neutral items. As this difference was absent in the rTMS condition when DLPFC was disrupted we obtained support for the view that DLPFC is involved in CIT type deceptive behavior and P300 is a sensitive signature of this. It must be stressed that compared to ERPs as a correlational measure of brain processes related to a specific behavior, TMS provided a means to exert a causal effect on the respective brain systems, thus strengthening the arguments in favor of DLPFC and P300 as the factors in CIT type deceptive behavior. The unexpected result was that no difference between left- and right-hemisphere stimulation was found, which did not support our specific TMS laterality related hypothesis. First, because contralateral homologous brain areas are strongly and reliably influenced by ipsilateral TMS (Rogasch and Fitzgerald, 2013), there may be a carryover effect so that right and left hemisphere manipulations become equivalent in certain specific conditions. Second, because in some of our other studies indicating especially the right DLPFC involvement in deception a different task compared to the CIT type of task used here was employed (Karton and Bachmann, 2011; Karton et al., 2014a), this difference may be a consequence of the different task demands and cognitive processes associated with these tasks. Indeed, as the most recent paper from our lab showed, when the context of the deceptive behavior was changed so as to increase motivation to lie, left hemisphere DLPFC manipulations became more effective in changing the rate of untruthful responses (Karton et al., 2014b).

The amount of correctly detected deceptions using the ERP methods analogous to the present one has been quite high in earlier studies, ranging between 85-95% (Rosenfeld et al., 2006, see also Rosenfeld et al., 1991; Farwell and Donchin, 1991; Allen et al., 1992). On the other hand, more recent studies have reported detection rates varying from 27% to 86% (Rosenfeld et al., 2004; Mertens and Allen, 2008; Abootalebi et al., 2009) or even no diagnostic power of the P300 at all (Gamer and Berti, 2010). When we used stricter criteria for ascertaining the level of individual effects and a modest number of trials with critical stimuli, deception could be detected reliably using P300 amplitude differences for only 22% of the subjects in our experiment. This is a result suggesting that rTMS based manipulations aimed at reducing sensitivity of psychophysiological markers to deception may not be universally applied. On the other hand, it is possible that different experimental / CIT-test setups using different types of stimuli are capable of increasing the sensitivity of the P300 based marker. (Actually, in one of our unpublished studies where instead of the written labels the photographs of the stolen items were used, the proportion of the subjects for whom P300 showed reliable effects of deception increased to 77%.) For example, it is true that the P300 response is stronger for more realistic stimulus material (Ambach et al., 2010; Cutmore et al., 2009). Abstract words used in our experiment may not have been optimal for guaranteeing very high sensitivity of the P300 in response to seeing a critical item. Another, more technical, aspect to consider is that P300 responses for critical stimuli depend on the number of irrelevant stimuli included in the experiment (Johnson and Rosenfeld, 1992; Hu et al., 2012). We did not have many items to be held in the working memory by the subjects. Therefore, subjects may have held not much irrelevant information in working memory which decreased the cognitive load during the task and made it easier to use countermeasures or produce P300 less conspicuous. Another limitation of our study concerns the unequal number of critical and other types of stimuli. This may have led to a stimulus probability related confounding effect on P300, which must be controlled in future studies of TMS effects on ERP markers of deception. But even with these reservations, we showed that rTMS can be used at least in some cases when one wants to reduce sensitivity of a subject to critical items in a CIT type lie-detection test.

Last but not least, as in practice the false-positive CIT results may have highly undesirable legal effects (e.g., Winograd and Rosenfeld, 2014), it is important to develop and use methods surpassing the 22% potential applicability of the P300 based CIT implied by the results of this study than to risk innocent accusation.

In one way or another, this study recommends that (i) P300 amplitude difference is used as the main EEG signature of deception, (ii) inter-individual variability of the susceptibility to the ERP based CIT should be acknowledged, (iii) in principle, pre-CIT rTMS can decrease the sensitivity of the testee’s brain to the test.

## Acknowledgements

This work was supported by Estonian Ministry of Education and Research, project SF0180027s12 (TSHPH0027), “Attention and Consciousness” and IUT20-40. We also thank Renate Rutiku, Jaan Aru, Carolina Murd and Anu Einberg for help they provided through various stages of this research.

## References

Abootalebi, V., Moradi, M. H., Khalilzadeh, M. A. (2009). A new approach for EEG feature extraction in P300-based lie detection. Computer Methods and Programs in Biomedicine, 94, 48–57. doi: 10.1016/j.cmpb.2008.10.001.

Allen, J J., Iacono, W. G., Danielson, K. D. (1992). The identification of concealed memories using the event-related potential and implicit behavioral measures: A methodology for prediction in the face of individual differences. Psychophysiology, 29, 504–522.

Ambach, W., Bursch, S., Stark, R., Vaitl, D. (2010). A Concealed Information Test with multimodal measurement. International Journal of Psychophysiology, 75, 258–267. doi: 10.1016/j.ijpsycho.2009.12.007.

Ambach, W., Dummel, S., Lüer, T., Vaitl, D. (2011). Physiological responses in a Concealed Information Test are determined interactively by encoding procedure and questioning format. International Journal of Psychophysiology, 81, 275–282. doi: 10.1016/j.ijpsycho.2011.07.010.

Bakeman, R. (2005). Recommended effect size statistics for repeated measures designs. Behavior Research Methods, 37(3), 379–384.

Berkman, E. T., Lieberman, M.D. (2010). Approaching the bad and avoiding the good: lateral prefrontal cortical asymmetry distinguishes between action and valence. Journal of Cognitive Neuroscience, 22, 1970–1979. doi: 10.1162/jocn.2009.21317.

Ben-Shakhar, G., Elaad, E. (2003). The validity of psychophysiological detection of information with the Guilty Knowledge Test: a meta-analytic review. Journal of Applied Psychology, 88(1), 131–151.

Carmel, D., Dayan, E., Naveh, A., Raveh, O., Ben-Shakhar, G. (2003). Estimating the validity of the Guilty Knowledge Test from simulated experiments: the external validity of mock crime studies. Journal of Experimental Psychology: Applied, 9(4), 261–269. doi: 10.1037/1076-898X.9.4.261.

Cho, S. S., Pellecchia, G., Ko, J. H., Ray, N., Obeso, I., Houle, S., Strafella, A. P. (2012). Effect of continuous theta burst stimulation of the right dorsolateral prefrontal cortex on cerebral blood flow changes during decision making. Brain Stimulation, 5, 116–123. doi: 10.1016/j.brs.2012.03.007.

Christ, S. E., Van Essen, D. C., Watson, J. M., Brubaker, L. E., McDermott, K. B. (2009). The contributions of prefrontal cortex and executive control to deception: evidence from activation likelihood estimate meta-analyses. Cereb. Cort. 19, 1557–1566. doi: 10.1093/cercor/bhn189

Cutmore, T. R., Djakovic, T., Kebbell, M. R., Shum, D. H. (2009). An object cue is more effective than a word in ERP-based detection of deception. Int. J. Psychophysiol. 71, 185–192. doi: 10.1016/j.ijpsycho.2008.08.003.

Eisenegger, C., Treyer, V., Fehr, E., Knoch, D. (2008). Time-course of “off-line” prefrontal rTMS effects--a PET study. NeuroImage 42(1), 379–384. doi: 10.1016/j.neuroimage.2008.04.172.

Farwell, L.-A. (2012). Brain fingerprinting: a comprehensive tutorial review of detection of concealed information with event-related brain potentials. Cogn. Neurodyn. 6, 115–154. doi: 10.1007/s11571-012-9192-2.

Farwell, L.-A., Donchin, E. (1991). The truth will out: Interrogative polygraphy (‘lie detector’) with event-related brain potentials. Psychophysiol. 28, 531–547.

Fertonani, A., Rosini, S., Cotelli, M., Rossini, P. M., Miniussi, C. (2010). Naming facilitation induced by transcranial direct current stimulation. Beh. Brain Res. 208, 311–318. doi: 10.1016/j.bbr.2009.10.030.

Gamer, M., Berti, S. (2010). Task relevance and recognition of concealed information have different influences on electrodermal activity and event-related brain potentials. Psychophysiol. 47, 355–364. doi: 10.1111/j.1469-8986.2009.00933.x.

Gamer, M., Bauermann, T., Stoeter, P., Vossel, G. (2007). Covariations among fMRI, skin conductance, and behavioral data during processing of concealed information. Hum. Brain Mapp. 28, 1287–1301. doi: 10.1002/hbm.20343.

Ganis, G., Keenan, J. P. (2009). The cognitive neuroscience of deception. Soc. Neurosci. 4, 465–472. doi: 10.1080/17470910802507660.

Ganis, G., Kosslyn, S. M., Stose, S., Thompson, W. I., Yurgelun-Todd, D. A. (2003). Neural correlates of different types of deception: An fMRI investigation. Cereb. Cort. 13, 830–836. doi: 10.1093/cercor/13.8.830.

Hallett, M. (2007). Transcranial magnetic stimulation: A primer. Neuron 55, 187–199. doi: http://dx.doi.org/10.1016/j.neuron.2007.06.026.

Hansenne, M., Laloyaux, O., Mardaga, S., Ansseau, M. (2004). Impact of low frequency transcranial magnetic stimulation on event-related brain potentials. Biol. Psychol. 67, 331–341.

Hortensius, R., Schutter, D. J. L., Harmon-Jones, E. (2012). When anger leads to aggression: induction of relative left frontal cortical activity with transcranial direct current stimulation increases the anger aggression relationship. SCAN 7, 342–347. doi: 10.1093/scan/nsr012.

Hu, X., Hegeman, D., Landry, E., Rosenfeld, J. P. (2012). Increasing the number of irrelevant stimuli increases ability to detect countermeasures to the P300-based Complex Trial Protocol for concealed information detection. Psychophysiol. 49, 85–95. doi: 10.1111/j.1469-8986.2011.01286.x.

Huffmeijer, R., Alink, L. R. A., Tops, M., Bakermans-Kranenburg, M. J., van IJzendoorn, M. H. (2012). Asymmetric frontal brain activity and parental rejection predict altruistic behavior: Moderation of oxytocin effects. Cogn. Aff. Beh. Neurosci. 12, 382–392. doi: 10.3758/s13415-011-0082-6.

Ito, A., Abe, N., Fujii, T., Hayashi, A., Ueno, A., Mugikura, S., Takahashi, S., Mori, E. (2012). The contribution of the dorsolateral prefrontal cortex to the preparation for deception and truth-telling. Brain Res. 1464, 43–52. doi: 10.1016/j.brainres.2012.05.004.

Jiang, W., Liu, H., Liao, J., Ma, X., Rong, P., Tang, Y., Wang, W. (2013). A functional MRI study of deception among offenders with antisocial personality disorders. Neurosci. 244, 90–98. doi: 10.1016/j.neuroscience.2013.03.055.

Johnson, M. M., Rosenfeld, J. P. (1992). Oddball-evoked P300-based method of deception detection in the laboratory II: utilization of non-selective activation of relevant knowledge. Int. J. Psychophysiol. 12(3), 289–306.

Jokinen, A., Santtila, P., Ravaja, N., Puttonen, S. (2006). Salience of guilty knowledge test items affects accuracy in realistic mock crimes. Int. J. Psychophysiol. 62(1), 175–184. doi: 10.1016/j.ijpsycho.2006.04.004

Karton, I., Bachmann, T. (2011). Effect of prefrontal transcranial magnetic stimulation on spontaneous truth-telling. Beh. Brain Res. 225, 209–214. doi: 10.1016/j.bbr.2011.07.028

Karton, I., Rinne, J. M., Bachmann, T. (2014a). Facilitating the right but not left DLPFC by TMS decreases truthfulness of object-naming responses. Beh. Brain Res. 271, 89–93. doi: 10.1016/j.bbr.2014.05.059.

Karton, I., Palu, A., Jõks, K., Bachmann, T. (2014b). Deception rate in a “lying game”: Different effects of excitatory repetitive transcranial magnetic stimulation of right and left dorsolateral prefrontal cortex not found with inhibitory stimulation. Neurosci. Lett. 583, 21–25.

Knoch, D., Fehr, E., 2007. Resisting the power of temptations: The right prefrontal cortex and self-control. Ann. New York Acad. Sci-es 1104, 123–134.

Kozel, F. A., Johnson, K. A., Grenesko, E. L., Laken, S. J., Kose, S., Lu, X., Pollina, D., et al. (2009). Functional MRI detection of deception after committing mock sabotage crime. J. Forens. Sci-es 54, 220–231. doi: 10.1111/j.1556-4029.2008.00927.x.

Kähkönen, S., Wilenius, J., Komssi, S., Ilmoniemi, R. J. (2004). Distinct differences in cortical reactivity of motor and prefrontal cortices to magnetic stimulation. Clin. Neurophysiol. 115, 583–588.

Langleben, D. D., Loughead, J. W., Bilker, W. B., Ruparel, K., Childress, A. R., Busch, S. I., et al. (2005). Telling truth from lie in individual subjects with fast event-related fMRI. Hum. Brain Mapp. 26, 262–272.

Luber, B., Fisher, C., Appelbaum, P. S., Ploesser, M., Lisanby, S. H. (2009). Noninvasive brain stimulation in the detection of deception: scientific challenges and ethical consequences. Beh. Sci-es & the Law, 27, 191–208. doi: 10.1002/bsl.860.

Luck, S.J. (2005). An Introduction to the Event-Related Potential Technique. Cambridge, MA: MIT Press.

Lykken, D. T. (1959). The GSR in the detection of guilt. Journal of Applied Psychology, 43, 385–388.

Lykken, D.T. (1979). The detection of deception. Psychol. Bull. 86, 47–53.

Mameli, F., Mrakic-Sposta, S., Vergari, M., Fumagalli, M., Macis, M., Ferrucci, R., Nordio, F., Consonni, D., Sartori, G., Priori, A. (2010). Dorsolateral prefrontal cortex specifically processes general - but not personal - knowledge deception: Multiple brain networks for lying *Beh*. Brain Res. 211, 164–168. doi:10.1016/j.bbr.2010.03.024.

Mertens, R., Allen, J. J. B. (2008). The role of psychophysiology in forensic assessments: Deception detection, ERPs, and virtual reality mock crime scenarios. Psychophysiol. 45, 286–298. DOI: 10.1111/j.1469-8986.2007.00615.x.

Nahari, G., Ben-Shakhar, G. (2011). Psychophysiological and behavioral measures for detecting concealed information: The role of memory for crime details. Psychophysiol. 48, 733–744. DOI: 10.1111/j.1469-8986.2010.01148.x.

Ott, D. V. M., Ullsperger, M., Jocham, G., Neumann, J., Klein, T. A. (2011). Continuous theta-burst stimulation (cTBS) over the lateral prefrontal cortex alters reinforcement learning bias. NeuroImage 57, 617–623. DOI: 10.1016/j.neuroimage.2011.04.038.

Polich, J. (2007). Updating P300: An integrative theory of P3a and P3b. Clin. Neurophysiol. 118, 2128–2148. doi: 10.1016/j.clinph.2007.04.019.

Priori. A., Mameli, F., Cogiamanian, F., Marceglia, S., Tiriticco, M., Mrakic-Sposta, S., Ferrucci, R., Zago, S., Polezzi. D., Sartori, G. (2008). Lie-specific involvement of dorsolateral prefrontal cortex in deception. Cereb. Cort. 18, 451–455.

Robertson, E. M., Thěoret, H., Pascual-Leone, A. (2003). Studies in cognition: the problems solved and created by transcranial magnetic stimulation. J. Cog. Neurosci. 15, 948–960.

Rogasch, N. C., Fitzgerald, P. B. (2013). Assessing cortical network properties using TMS–EEG. Hum. Brain Mapp. 34, 1652–1669. doi: 10.1002/hbm.22016.

Rosenfeld, J. P., Angell, A., Johnson, M., Qian, J.-H. (1991). An ERP-based, control-question lie detector analog: Algorithms for discriminating effects within individuals’ average waveforms. Psychophysiol. 28, 319–335.

Rosenfeld, J. P., Hu, X., Labkovsky, E., Meixner, J., Winograd, M. R. (2013). Review of recent studies and issues regarding the P300-based complex trial protocol for detection of concealed information. Int. J. Psychophysiol. 90, 118–134. doi: 10.1016/j.ijpsycho.2013.08.012.

Rosenfeld, J. P. (2011). P300 in detecting concealed information. In B. Verschuere, G. Ben-Shakhar, & E. Meijer (Eds.), Memory Detection: Theory and Application of the Concealed Information Test (pp. 63–89). Cambridge: Cambridge University Press.

Rosenfeld, J. P., Soskins, M., Bosh, G., Ryan, A. (2004). Simple effective countermeasures to P300-based tests of detection of concealed information. Psychophysiol. 41, 205–219.

Rosenfeld, J. P., Labkovsky, E. (2010). New P300-based protocol to detect concealed information: Resistance to mental countermeasures against only half the irrelevant stimuli and a possible ERP indicator of countermeasures. Psychophysiol. 47, 1002–1010. doi: 10.1111/j.1469-8986.2010.01024.x.

Rosenfeld, J. P., Biroschak, J. R., Furedy, J. J. (2006). P300-based detection of concealed autobiographical versus incidentally acquired information in target and non-target paradigms. Int. J. Psychophysiol. 60(3), 251–259.

Shackman, A. J., McMenamin, B. W., Maxwell, J. S., Greischar, L. L., Davidson, R. J. (2009). Right dorsolateral prefrontal cortical activity and behavioral inhibition. Psychol. Sci. 20, 1500–1506. doi: 10.1111/j.1467-9280.2009.02476.x.

Shafi, M. M., Westover, M. B., Fox, M. D., Pascual-Leone, A. (2012). Exploration and modulation of brain network interactions with noninvasive brain stimulation in combination with neuroimaging. Eur. J. Neurosci. 35, 805–825. doi: 10.1111/j.1460-9568.2012.08035.x.

Soskins, M., Rosenfeld, J. P., Niendam, T. (2001). The case for peak to-peak measurement of P300 recorded at .3 Hz high pass filter settings in detection of deception. Int. J. Psychophysiol. 40, 173–180.

Steinbeis, N., Bernhardt, B. C., Singer, T. (2012). Impulse control and underlying functions of the left DLPFC mediate age-related and age-independent individual differences in strategic social behavior. Neuron 73, 1040–1051. doi: 10.1016/j.neuron.2011.12.027.

Tassy, S., Oullier, O., Duclos, Y., Coulon, O., Mancini, J., Deruelle, C., Attarian, S., Felician, O., Wicker, B. (2012). Disrupting the right prefrontal cortex alters moral judgment. SCAN 7, 282–288. doi: 10.1093/scan/nsr008.

Thut, G., Pascual-Leone, A., 2010. A review of combined TMS-EEG studies to characterize lasting effects of repetitive TMS and assess their usefulness in cognitive and clinical neuroscience. Brain Topog. 22, 219–232. doi: 10.1007/s10548-009-0115-4.

Torii, T., Sato, A., Iwahashi, M., Iramina, K. (2012). Transition of After Effect on P300 by Short-Term rTMS to Prefrontal Cortex. IEEE Trans. Magn. 48(11), 2873–2876. doi: 10.1109/TMAG.2012.2204432

Verschuere, B., Ben-Shakhar, G., Meijer, E. (Eds.). (2011). Memory Detection: Theory and Application of the Concealed Information Test. Cambridge: Cambridge University Press.

Winograd, M. R., Rosenfeld, J. P. (2014). The impact of prior knowledge from participant instructions in a mock crime P300 Concealed Information Test. Int. J. Psychophysiol. 94(3), 473–481. http://dx.doi.org/10.1016/j.ijpsycho.2014.08.002.

